# Population genomic structure and genome-wide linkage disequilibrium in farmed Atlantic salmon (*Salmo salar* L.) using dense SNP genotypes

**DOI:** 10.1101/406405

**Authors:** Agustin Barria, Maria E. López, Grazyella Yoshida, Roberto Carvalheiro, José M. Yáñez

## Abstract

Chilean Farmed Atlantic salmon (*Salmo salar*) populations were established with individuals of both European and North American origins. These populations are expected to be highly genetically differentiated due to evolutionary history and poor gene flow between ancestral populations from different continents. The extent and decay of linkage disequilibrium (LD) among single nucleotide polymorphism (SNP) impacts the implementation of genome-wide association studies and genomic selection and provides relevant information about demographic processes of fish populations. We assessed the population structure and characterized the extent and decay of LD in three Chilean commercial populations of Atlantic salmon with North American (NAM), Scottish (SCO) and Norwegian (NOR) origin. A total of 151 animals were genotyped using a 159K SNP Axiom® myDesign™ Genotyping Array. A total of 40K, 113K and 136 K SNP markers were used for NAM, SCO and NOR populations, respectively. The principal component analysis explained 86.7% of the genetic diversity between populations, clearly discriminating between populations of North American and European origin, and also between European populations. Admixture analysis showed that the Scottish and North American populations likely come from one ancestral population, while the Norwegian population probably originated from more than one. NAM had the lowest effective size, followed by SCO and NOR. Large differences in the LD decay were observed between populations of North American and European origin. A r^2^ threshold of 0.2 was estimated for marker pairs separated by 8,000 Kb, 42 and 64 Kb in the NAM, NOR and SCO populations, respectively. In this study we show that this SNP panel can be used to detect association between markers and traits of interests and also to capture high-resolution information for genome-enabled predictions. Also, we suggest the feasibility to achieve higher prediction accuracies by using a small SNP data set as was used with the NAM population.

## Background

Atlantic salmon (*Salmo salar*) is one of the species of farmed fish with the highest commercial value in aquaculture (FAO, 2016b). Chile is the second largest producer, generating nearly 532,000 tons of fish in 2016 (FAO, 2016a). All of the Atlantic salmon populations farmed in Chile were introduced from three main geographical origins i) North America, ii) Scotland and iii) Norway. These populations also represent the main origins of cultured Atlantic salmon worldwide. Breeding programs for Atlantic salmon were first established in Norway during the early 1970´s (Gjedrem et al., 2012). Since then, there has been an increased interest in implementing genetic improvement programs for salmon in the most important producer countries, including Australia, Chile, Iceland, Ireland, Scotland and Norway. The main traits included in the breeding objectives of Atlantic salmon are growth, disease resistance, carcass quality and age at sexual maturation (Rye et al., 2010).

Recent advances in next-generation sequencing and high-throughput genotyping technologies have allowed the development of valuable genomic resources in aquaculture species (Yáñez et al., 2015). For instance, dense single nucleotide polymorphism (SNP) panels have been developed for Atlantic salmon (Houston et al., 2014; Yáñez et al., 2016). Genetic evaluations for traits that are difficult to measure in selection candidates, such as disease resistance and carcass quality traits, can be more accurate when integrating genome-wide SNP information, in what has been called genomic selection (Meuwissen et al., 2001; Sonesson and Meuwissen, 2009). Genomic selection exploits the linkage disequilibrium (LD) that exists between SNP and quantitative trait loci (QTL) or causative mutations that are involved in the variation of the trait (Goddard and Hayes, 2009), increasing the accuracy of genome-enabled estimated breeding values (GEBVs) in farmed salmon species (Bangera et al., 2017; Barría et al., 2018; Ødegård et al., 2014; Tsai et al., 2016; Vallejo et al., 2017; Yoshida et al., 2018). Furthermore, association mapping through genome wide association studies (GWAs) is a useful approach to detect genomic regions and genes involved in economically important traits for salmon aquaculture and they also rely on LD between the QTL and SNP markers. Thus an adequate SNP density is required to assure that all QTL are in LD with a marker (Flint-Garcia et al., 2003).

In addition, knowing the extent and pattern of LD can be used to help explore different evolutionary forces that may affect certain regions of the genome (Ardlie et al., 2002). Because it is affected by population growth, genetic drift, admixture or migration, population structure, variable recombination rates and artificial/natural selection LD can be variable among populations and loci (Ardlie et al., 2002). Different measures of LD between two loci have been proposed, among them the absolute value of *D’* (also called Lewontin’s *D*’) and r^2^ are the most widely used. *D’* = 1, indicates no recombination between loci and complete LD, while values less than 1 indicate that loci have been separated by recombination, but there is no clear interpretation. D’ estimations are overestimated in small sample sizes and low frequencies of minor allele, therefore, high values of D’ can be obtained even when markers are in linkage equilibrium (Ardlie 2002). Therefore, r^2^, the squared correlation between alleles at two loci, is the most accepted measure for comparing and quantifying LD (Pritchard and Przeworski, 2001). r^2^ =1, only if two markers have not been separated by recombination and have the same allele frequency, showing perfect linkage disequilibrium, one marker provides complete information about the other marker (Ardlie 2002).

To date, several studies have been performed to determine the levels and extent of LD in livestock species such as dairy (Bohmanova et al., 2010; Khatkar et al., 2008), beef cattle (Espigolan et al., 2013; Lu et al., 2012; McKay et al., 2007; Porto-Neto et al., 2014), pigs (Ai et al., 2013; Badke et al., 2012), goats (Mdladla et al., 2016; Visser et al., 2016) and sheep (Prieur et al., 2017). Moreover, some studies have related patterns of LD with genomic regions subjected to selection in domestic species (Prasad et al., 2008). Recent studies have also aimed at characterizing the levels of LD in farmed aquaculture species, such as rainbow trout (Rexroad and Vallejo, 2009; Vallejo et al., 2018), coho salmon (Barria et al., 2018) and Atlantic salmon (Kijas et al., 2017). However, until now there have been no comprehensive studies aiming at characterizing and comparing levels and extent of LD in commercial Atlantic salmon populations that include the three main geographical origins. The goal of this study was to (a) assess the levels of LD in farmed Atlantic salmon populations with three different geographical origins (i.e. Canada, Scotland and Norway); (b) calculate the effective population size for each breeding population and (c) estimate the population structure and genetic admixture of each population.

## Methods

### Populations and samples

The current study is comprised of 151 Atlantic salmon individuals from three different commercial populations cultivated in the South of Chile, that have different geographical origins. These fish were obtained from Chilean farmed populations, which were originated from imported stocks. The Norwegian population was comprised of 71 fish belonging to a breeding population derived from the Mowi strain, which was established in the late 1960s using fish from west coast rivers in Norway (NOR), River Bolstad in the Vosso watercourse, River Årøy and Maurangerfjord area (Verspoor et al., 2007). Ova of this strain were introduced into Ireland from 1982 to 1986 (Norris et al., 1999) and from there, they were imported to Chile for farming purposes in the 1990´s (Solar, 2009). Since 1997 this population has been selected for rapid growth in Chile (Correa et al., 2015, 2017, Yáñez et al., 2013, 2014). A second population of 43 fish of Scottish origin (SCO) was comprised of samples from a strain derived from fish from Loch Lochy, located on the West Coast of Scotland. Fish of this strain are described as a stock with rapid growth potential and a high early maturation grilsing rate (Johnston et al., 2000). During the 1980´s, eggs from the Scottish population were introduced to Chile to establish an aquaculture broodstock. The third population used in this study was comprised of 37 fish of North American (NAM) origin; belonging to a domestic strain established in the 1950´s, using ova from the Gaspé Bay (Québec, Canada). It is presumed that fish of this strain were transferred and kept at an aquaculture hatchery located in the state of Washington, USA for two generations. Fertilized eggs of this strain were introduced from Washington to Chile between 1996 and 1998 (López et al., 2018).

### Genotyping

Fin clip samples from individuals from the three populations were obtained for genomic DNA extraction and further genotyping. Genotyping was carried out using a 200K Affymetrix Axiom ® myDesign Custom Array as described by (Yáñez et al., 2016). This dense SNP array contains 151,509 polymorphic SNPs with unique position and evenly distributed markers across the genome. A total of 2,302 (1.6 %) SNPs were discarded prior to analysis due to unknown chromosomal location on the *Salmo salar* reference genome (Yáñez et al., 2016). Quality control of genotypes was performed using Axiom Genotyping Console (AGT, Affymetrix) and SNPolisher for R, according to the Best Practices procedures indicated by the array manufacturer (http://media.affymetrix.com/support/downloads/manuals/axiom_best_practice_supplement_user_guide.pdf). All subsequent analyses were assessed using a different subset of SNPs. Markers and sample quality control (QC) were performed using PLINK software v1.09 (Purcell et al., 2007).

### Estimation of LD

Linkage disequilibrium as Pearson’s squared correlation coefficient (r^2^) was chosen over |D’| to predict the LD between each pair of molecular markers. This statistic is less sensitive to bias caused by differences in allelic frequencies (Ardlie et al., 2002), more appropriate for biallelic markers (Zhao et al., 2005) and can be used to compare the results with previous studies in salmonid species and other domestic animals. Genotypes were coded as 2, 1 and 0 in function of the number of non-reference alleles. The pair-wise LD as r^2^ was calculated for each population and within chromosomes using Plink v1.09 using the formula proposed by Hill and Robertson (Hill and Robertson, 1968). For each SNP pair, bins of 100kb were created based on pairwise physical distance. The extent and decay of the LD, was visualized by plotting the average r^2^ within each bin from 0 up to 10 Mb, using R software (R Core Team, 2016). Quality control was assessed separately for each population. SNPs with minor allele frequency (MAF) lower than 5%, significantly deviated from Hardy-Weinberg Equilibrium (HWE) (p < 1e-6), and a call rate of SNPs lower than 95% were excluded. Samples missing more than 5% of the genotype were also excluded.

### Effective population size

Historical effective population size (Ne) was estimated using SNeP v1.1 (Barbato et al., 2015). SNeP software estimates Ne using LD data calculated through the following formula proposed by Corbin et al. (Corbin et al., 2012):

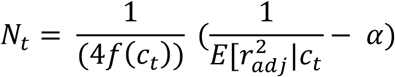

Where N_t_ is the effective population size t generations ago, c_t_ is the recombination rate t generations ago, r^2^adj is the estimated LD adjusted for sampling bias and *α* is a constant. Recombination rate was calculated as suggested by Sved (Sved, 1971):

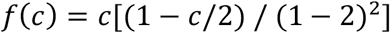

The minimum and maximum distance used between SNPs for Ne estimation was 0 and 5 Mb, respectively. Data was grouped in 30 distance bins of 50 Kb each. Finally, Ne was estimated from the r^2^ values calculated for the mean distance of each distance bin. The SNP and sample data set used for Ne estimation was the same as previously used for LD estimation.

### Population structure and genetic admixture analysis

To investigate genetic structure among populations, we performed a principal component analysis (PCA) implemented in PLINK v1.09. Visualization of the first two PCA were plotted along two axes in R. Additionally, we performed a model-based clustering using ADMIXTURE v.1.3 (Alexander et al., 2009) to investigate individual admixture proportions. We tested several K, and selected the optimum K value according to the lowest value of the cross-validation error. PCA analysis was assessed using common SNPs between the three populations after the previous QC, which was done separately for each population. For the admixture analysis, raw data for the three populations was merged and QC was performed. However, unlike the QC carried out previously, no MAF filter was done.

## RESULTS

### SNP quality control

No animals from the three populations were removed after quality control, giving genotype data from 151 individuals (37, 43 and 71 from NAM, SCO and NOR, respectively). A total of 40,316 (27.02%), 113,282 (75.92%) and 136,437 (91.44%) SNP markers passed the QC criteria for the NAM, SCO and NOR populations, respectively. Filtered SNPs differed significantly between populations of North American or European origin. 106 K SNPs were excluded from the NAM population by a low MAF, representing 70% of the total available markers in the array. The markers excluded by MAF in SCO and NOR populations reached 23 and 7.3%, respectively. A summary of the number of fish genotyped from each population, number of SNPs excluded by HWE, MAF and the final number of SNPs per population are shown in Table 1.

**Table 1.** Summary of results from quality control of SNPs for each farmed population genotyped with the 200K SNP Array

| Population origin <sup>1</sup> | **n<sub>g</sub>** <sup>2</sup> | Callrate <sup>3</sup> | HWE <sup>4</sup> | MAF < 0.05 <sup>5</sup> | Final number of SNPs <sup>6</sup> |
| --- | --- | --- | --- | --- | --- |
| NAM | **37** | **2600** | **188** | **106103** | **40316** |
| SCO | **43** | **923** | **62** | **34940** | **113282** |
| NOR | **71** | **1159** | **128** | **11083** | **136437** |

<sup>1</sup> Geographical origin of each farmed population: NAM = North America; SCO = Scotland; NOR = Norway.
<sup>2</sup> Number of genotyped samples
<sup>3</sup> Number of Excluded SNPs by the genotype call rate < 0.95
<sup>4</sup> Number of excluded SNPs by Hardy–Weinberg Equilibrium.
<sup>5</sup> Number of excluded SNPs with minor allele frequency < 0.05.
<sup>6</sup> Number of SNPs used in the subsequent analysis for each population.

### Summary statistics for each population

Summary statistics of each chromosome’s length and LD estimates among SNPs for each chromosome and population are shown in Table 2. The markers spanned 2,218.6 Mb, 2,233.2 Mb and 2,235.3 Mb of the genome for the NAM, SCO and NOR populations, respectively. Average r^2^ between adjacent SNPs reach up to 0.35 ± 0.35 in the NAM population. These values were lower for SCO and NOR populations (0.13 ± 0.16 and 0.08 ± 0.12, respectively). The average LD, measured as r^2^, between adjacent markers across the 29 chromosomes, ranged from 0.16 to 0.35, 0.08 to 0.13 and from 0.05 to 0.08 in NAM, SCO and NOR populations, respectively (Table 2). These results indicate that average levels of LD among syntenic SNPs are considerably higher in both populations with European origin. Also, LD for the SCO population is slightly higher when compared with the NOR population.

**Table 2.** Estimated chromosome length and average linkage disequilibrium values for three Chilean farmed populations of Atlantic salmon with Norwegian (NOR), Scottish (SCO) and North American (NAM) origin.

| Chr<sup>a</sup> | NAM |  | SCO |  | NOR |  |
| --- | --- | --- | --- | --- | --- | --- |
|  | Length (Mb) | Average r<sup>2</sup> | Length (Mb) | Average r<sup>2</sup> | Length (Mb) | Average r<sup>2</sup> |
| 1 | 158.79 | 0.32 ± 0.32 | 158.94 | 0.11 ± 0.14 | 158.94 | 0.06 ± 0.09 |
| 2 | 72.66 | 0.16 ± 0.20 | 72.90 | 0.08 ± 0.11 | 72.92 | 0.05 ± 0.08 |
| 3 | 91.90 | 0.31 ± 0.33 | 92.18 | 0.08 ± 0.11 | 92.31 | 0.05 ± 0.07 |
| 4 | 80.75 | 0.23 ± 0.27 | 81.73 | 0.13 ± 0.16 | 82.37 | 0.06 ± 0.09 |
| 5 | 78.04 | 0.21 ± 0.25 | 80.04 | 0.08 ± 0.12 | 80.42 | 0.05 ± 0.08 |
| 6 | 86.67 | 0.21 ± 0.25 | 86.81 | 0.08 ± 0.11 | 86.87 | 0.05 ± 0.08 |
| 7 | 56.13 | 0.20 ± 0.23 | 58.39 | 0.08 ± 0.11 | 58.39 | 0.05 ± 0.07 |
| 8 | 23.98 | 0.35 ± 0.35 | 25.17 | 0.08 ± 0.11 | 25.51 | 0.05 ± 0.08 |
| 9 | 141.54 | 0.33 ± 0.31 | 141.70 | 0.11 ± 0.14 | 141.70 | 0.05 ± 0.08 |
| 10 | 116.05 | 0.22 ± 0.26 | 116.06 | 0.10 ± 0.13 | 116.10 | 0.08 ± 0.12 |
| 11 | 93.49 | 0.27 ± 0.28 | 93.81 | 0.10 ± 0.14 | 93.84 | 0.06 ± 0.09 |
| 12 | 91.01 | 0.21 ± 0.24 | 91.82 | 0.12 ± 0.15 | 91.82 | 0.06 ± 0.09 |
| 13 | 107.41 | 0.30 ± 0.30 | 107.71 | 0.12 ± 0.15 | 107.71 | 0.08 ± 0.11 |
| 14 | 93.61 | 0.23 ± 0.28 | 93.62 | 0.09 ± 0.13 | 93.81 | 0.07 ± 0.11 |
| 15 | 103.20 | 0.30 ± 0.31 | 103.81 | 0.12 ± 0.15 | 103.81 | 0.07 ± 0.10 |
| 16 | 87.43 | 0.25 ± 0.28 | 87.70 | 0.11 ± 0.14 | 87.70 | 0.06 ± 0.09 |
| 17 | 57.08 | 0.24 ± 0.26 | 57.44 | 0.10 ± 0.13 | 57.44 | 0.05 ± 0.08 |
| 18 | 69.96 | 0.32 ± 0.31 | 70.44 | 0.13 ± 0.17 | 70.44 | 0.05 ± 0.08 |
| 19 | 82.01 | 0.22 ± 0.26 | 82.83 | 0.09 ± 0.12 | 82.83 | 0.07 ± 0.11 |
| 20 | 86.02 | 0.32 ± 0.31 | 86.64 | 0.12 ± 0.15 | 86.64 | 0.06 ± 0.10 |
| 21 | 57.47 | 0.26 ± 0.29 | 57.93 | 0.08 ± 0.11 | 57.93 | 0.05 ± 0.08 |
| 22 | 62.97 | 0.22 ± 0.26 | 63.31 | 0.09 ± 0.12 | 63.31 | 0.06 ± 0.09 |
| 23 | 49.59 | 0.35 ± 0.33 | 49.71 | 0.10 ± 0.13 | 49.71 | 0.05 ± 0.08 |
| 24 | 47.65 | 0.28 ± 0.29 | 48.13 | 0.11 ± 0.15 | 48.13 | 0.05 ± 0.08 |
| 25 | 51.27 | 0.26 ± 0.28 | 51.43 | 0.12 ± 0.16 | 51.43 | 0.06 ± 0.09 |
| 26 | 47.45 | 0.22 ± 0.25 | 47.75 | 0.11 ± 0.15 | 47.75 | 0.07 ± 0.10 |
| 27 | 43.45 | 0.22 ± 0.27 | 43.73 | 0.11 ± 0.15 | 43.77 | 0.05 ± 0.08 |
| 28 | 38.60 | 0.24 ± 0.26 | 39.04 | 0.08 ± 0.12 | 39.25 | 0.05 ± 0.08 |
| 29 | 42.41 | 0.17 ± 0.22 | 42.38 | 0.08 ± 0.11 | 42.41 | 0.05 ± 0.07 |

<sup>a</sup> Chromosome

Average SNP density per chromosome per Mb ranged from 10.28 to 24.39, 36.23 to 63.61 and from 45.13 to 71.92 for the NAM, SCO and NOR populations, respectively (**Table S1**). These differences are explained by the number of SNPs used within each population. However, the markers are uniformly distributed along the 29 chromosomes. All three populations showed a similar mean MAF of 0.26 ± 0.13, 0.28 ± 0.13 and 0.29 ± 0.13 for the NAM, SCO and NOR, respectively. The mean MAF per chromosome ranged from 0.28 to 0.30 in the NAM population. For both populations with European origin, the MAF ranged from 0.22 to 0.29. Except for the MAF values between 0.05 and 0.09, the proportion of loci ranged from 0.18 to 0.27 among the three populations (Figure 1).

**Figure 1.**
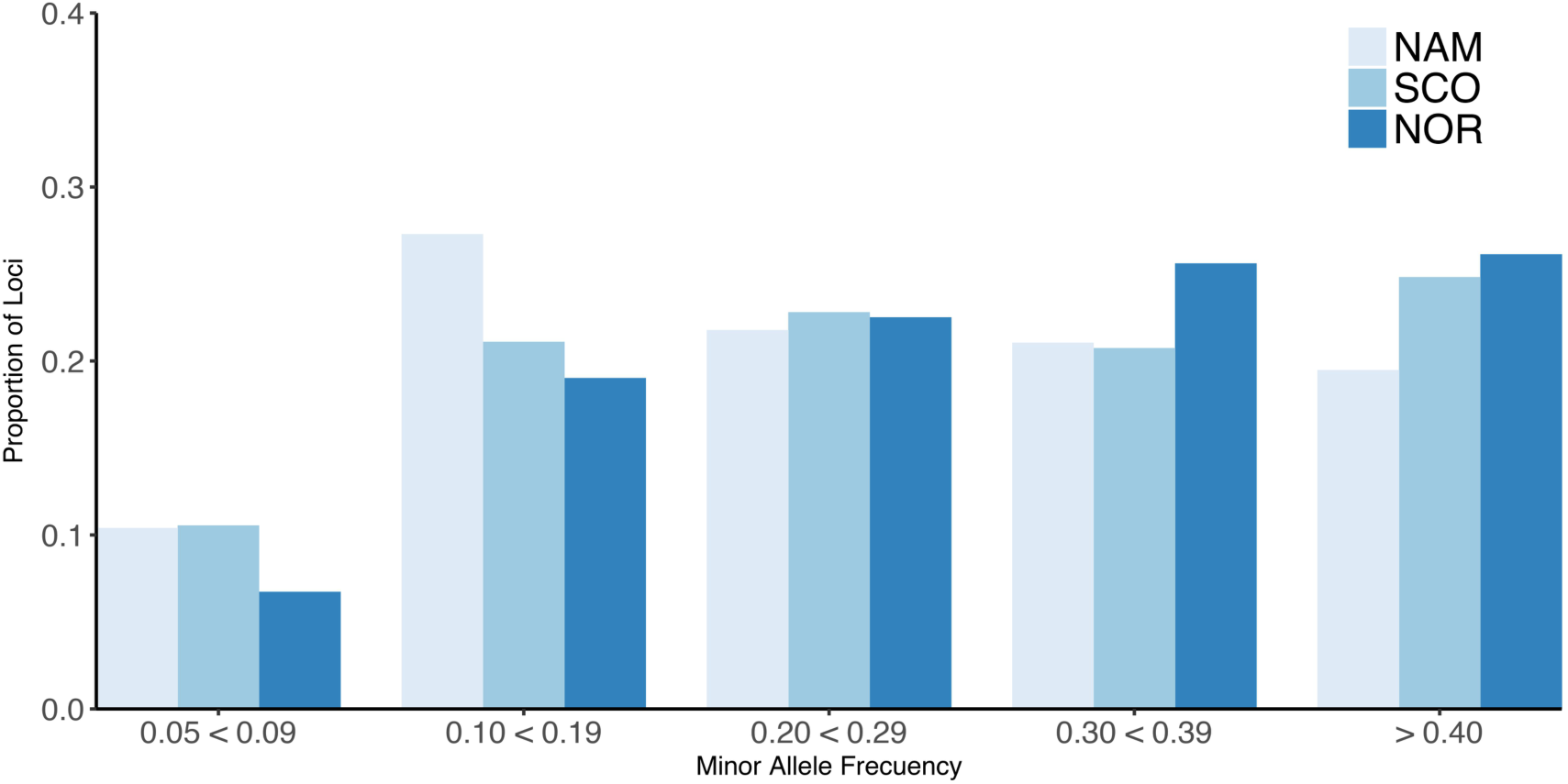
Minor allele frequency (MAF) for farmed Atlantic salmon populations. Proportion of loci for different values of Minor allele frequency for three Chilean farmed Atlantic salmon populations with different origin. North America (NAM), Scottish (SCO) and Norwegian (NOR).

### Population structure and genetic admixture

A subset of 33,124 SNPs with MAF > 5% and under Hardy-Weinberg equilibrium (HWE) shared among the three populations were used for principal component analysis (PCA) (Figure 2). Principal components 1 and 2 together accounted for 86.7% of the total genetic variation. These components clearly revealed three different clusters, corresponding to the Atlantic salmon with North American (NAM), Scottish (SCO) and Norwegian (NOR) origin. The first principal component discriminates populations with North American and European origin and accounted for 58.4% of the total variation. The second principal component accounted for 28.3% of the total variance and divided the two European populations into two clusters, corresponding to Scottish and Norwegian populations respectively.

**Figure 2.**
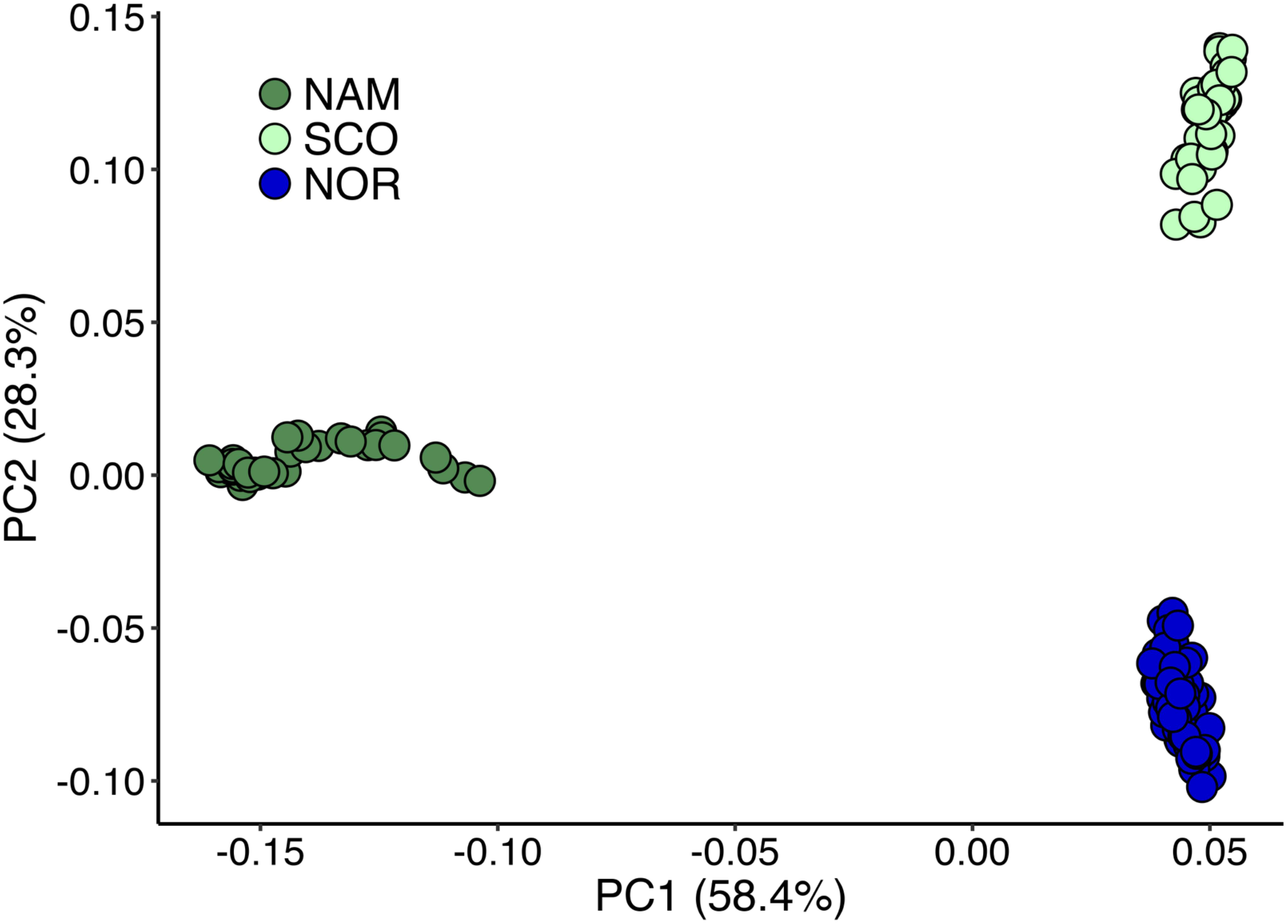
Genetic diversity of Atlantic salmon populations revealed by principal component analysis. Principal component analysis for three Chilean Atlantic salmon breeding populations with different geographical origin. North America (NAM), Scottish (SCO) and Norwegian (NOR).

Similarly to the results form PCA, ADMIXTURE at K=3, separated the three populations according to their geographical origins, however, based on the lowest cross-validations error, *K* = 4 was identified as the optimal number of ancestral populations; indicating a small European component in the North American population, as well as a Scottish component in the Norwegian population and vice versa. This analysis also revealed a high level of admixture in the Norwegian population, showing two ancestral groups. (Figure 3)

**Figure 3.**
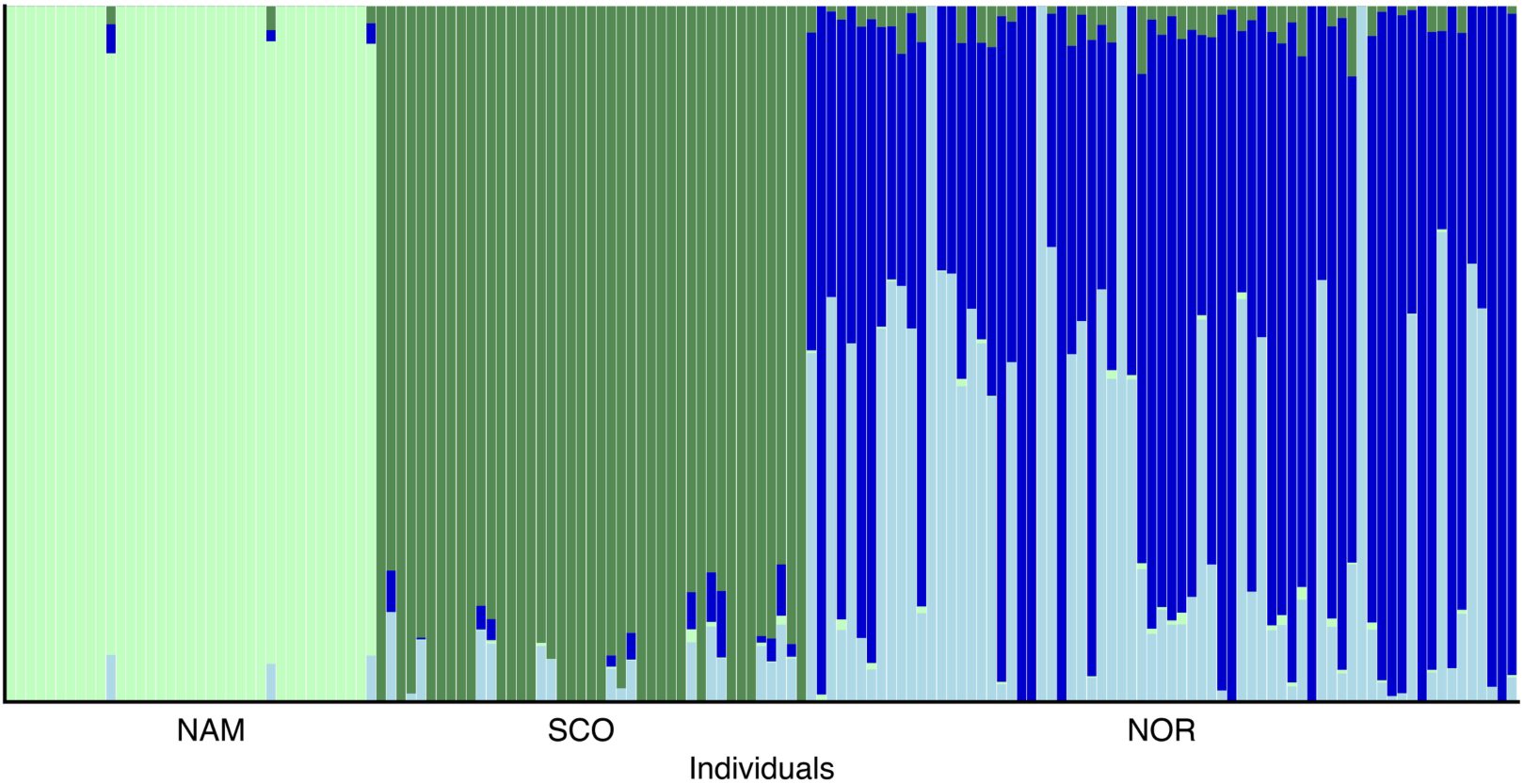
Population structure (k = 4) in three farmed populations of Atlantic salmon. Population structure in three farmed populations of Atlantic salmon with North America (NAM), Scottish (SCO) and Norwegian (NOR) origin. Each vertical line represents an individual, and each color a different theoretical ancestral population.

### Linkage disequilibrium analyses

The LD decay was estimated for each population as a function of physical distance. SNP pairs were sorted in 100 kb-bins based on the distance between pairs. Average r^2^ values were estimated for each bin. As estimated in other domestic animals (Badke et al., 2012; Barria et al., 2018; Kijas et al., 2017; Makina et al., 2015), genome-wide average LD declines with increasing physical distance between markers. Figure 4 shows an overview of the decay of r^2^ as a function of distance for each population. A slow decay was observed for the NAM population, while the decay was faster in both populations with European origin. The average distance at which the LD value reached 0.2, varied between populations. For the NAM population the distance reached ∼ 8,000 Kb. For the SCO and NOR populations, the distance decreased drastically, corresponding to ∼ 64 and 42 Kb, respectively. Average r^2^ for the first bin at distances of 0.5, 1.0, 5.0 and 10.0 Mb are shown in Table 3. Mean r^2^ within the first bin was larger for the NAM population (r^2^ = 0.62), followed by the SCO and NOR (r^2^ = 0.29 and 0.25, respectively). Average r^2^ for SNP pairs with a mean distance of 1.0 Mb was 0.31, 0.12 and 0.07 for the NAM, SCO and NOR populations, respectively. These values decreased to 0.19, 0.08 and 0.05, when average distance between SNPs reached up to 10.0 Mb for NAM, SCO and NOR, respectively.

**Table 3.** Mean linkage disequilibrium (*r*^2^) at different distances in three Chilean farmed populations of Atlantic salmon with North American (NAM), Scottish (SCO) and Norwegian (NOR) origin.

| **Population** | **0.1 Mb** | **0.5 Mb** | **1.0 Mb** | **5.0 Mb** | **10.0 Mb** |
| --- | --- | --- | --- | --- | --- |
| **NAM** | 0.62 | 0.36 | 0.31 | 0.22 | 0.19 |
| **SCO** | 0.29 | 0.13 | 0.12 | 0.09 | 0.08 |
| **NOR** | 0.25 | 0.08 | 0.07 | 0.05 | 0.05 |

**Figure 4.**
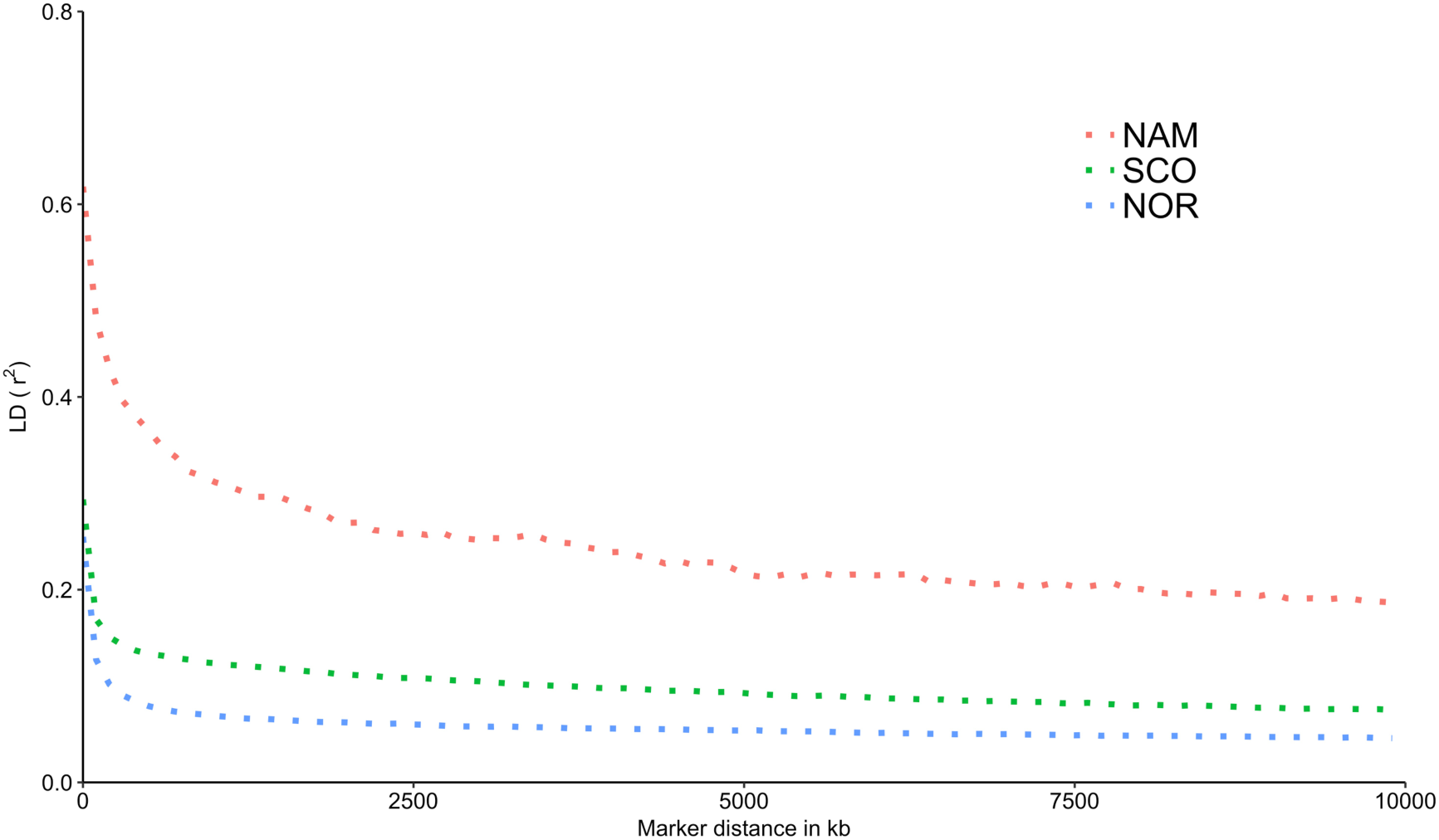
Decay of average linkage disequilibrium (*r*^2^) over distance in three farmed populations of Atlantic salmon. Average linkage disequilibrium between markers measured as *r*^2^ in three farmed populations of Atlantic salmon with North America (NAM), Scottish (SCO) and Norwegian (NOR) origin.

### Effective population size

Estimated effective population size differed among populations. Figure 5 shows the trends in Ne up to 85 (Figure 5A) and 1516 (**Figure 5B**) generations ago, respectively. Within this range of generations, Atlantic salmon with NAM origin had the smallest Ne, followed by SCO and NOR populations. These Ne values ranged from 19 to 1,325; 48 to 2,383 and from 86 to 2,717 for NAM, SCO and NOR populations, respectively.

**Figure 5.**
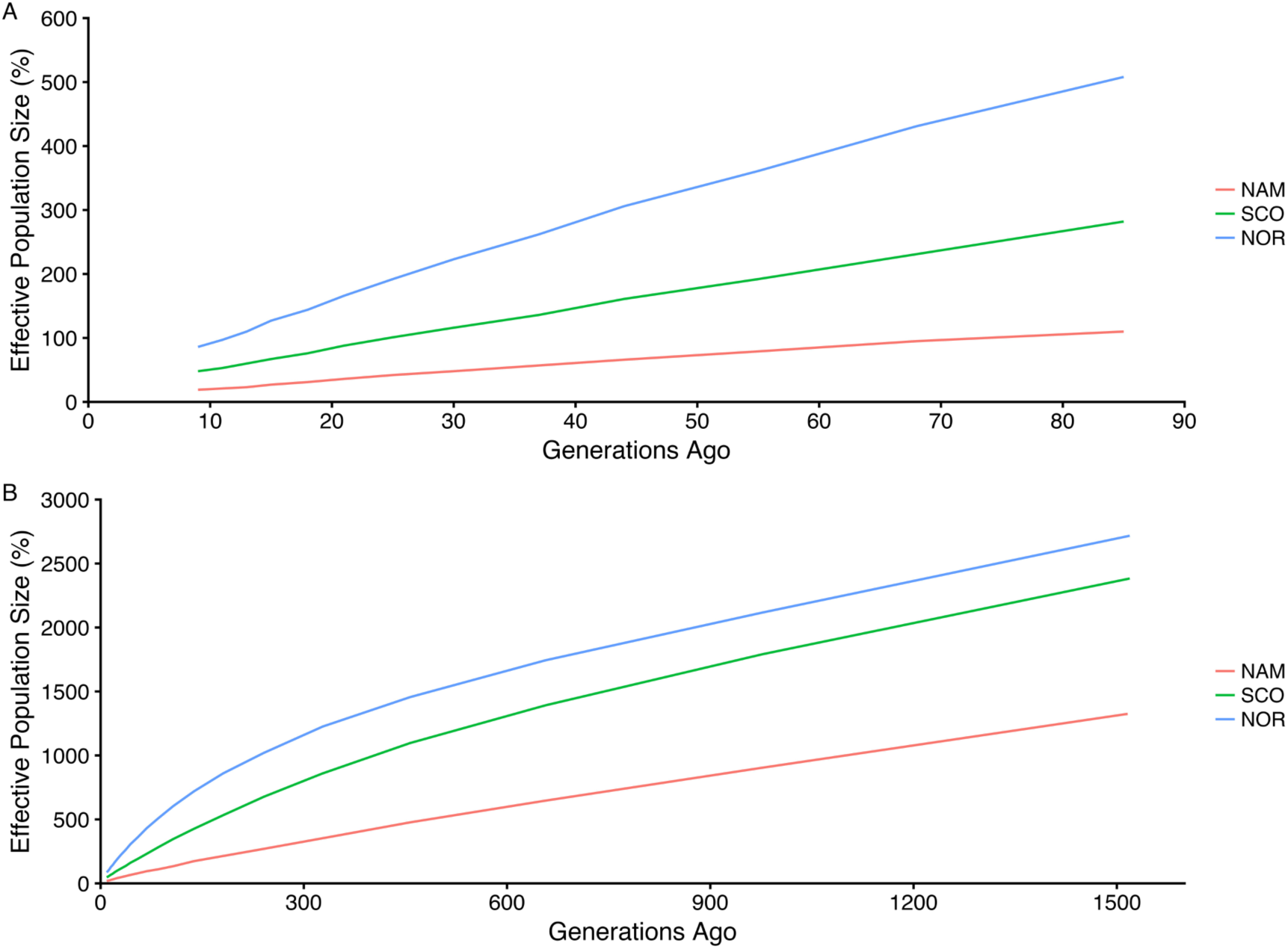
Estimated effective population size in three farmed populations of Atlantic salmon. Historic effective population size from 9 up to 85 (A) and 1516 (B) generations ago for three Atlantic salmon farmed populations with North America (NAM), Scottish (SCO) and Norwegian (NOR) origin.

## DISCUSSION

The study of extent and decay of whole-genome LD can aid in the understanding of demographic processes experienced by populations. Processes such as founder effect, admixture and genetic drift in conjunction with recombination and mutations are key elements determining LD. Similarly, other factors that affect LD include inbreeding, admixture and selection (Gaut and Long, 2003); which has resulted in studies aimed at estimating LD variation between populations (Ai et al., 2013; Al-mamun et al., 2015; Mdladla et al., 2016; Yang et al., 2014).

This is the first study aimed at characterizing the decay and extent of LD in three different Atlantic salmon breeding populations established in Chile, representing the three main geographical origins of this cultured species now in Chile. All three populations have been subjected to artificial selection for growth related traits. Individuals used in this study were selected having non-common ancestors for three generations back to avoid inflated LD estimations that are likely to occur due to high kinship relationships (Gutierrez et al., 2015).

The results presented here indicate the existence of differential average levels of LD across the genome between these Atlantic salmon populations. Large differences between North American and European populations were expected considering that they belong to two different lineages probably separated by more than 1,000,000 years (Rougemont and Bernatchez, 2018). The highest level of LD was in the NAM population and could reflect demographic process in its strain formation, as well as demographic events in its wild progenitors. It is well known that North American populations of Atlantic salmon have lower genetic diversity than European populations (Bourret et al., 2013; Makinen et al., 2014), as migrations probably favored only a few individuals colonizing North America, reducing effective sizes and causing a major effect of genetic drift (Rougemont and Bernatchez, 2018). Additionally, artificial selection in this population has been stronger than in SCO, which has been undergoing genetic improvement using mass selection. On the other hand, the lowest level of LD found in NOR is consistent with its admixed origin (Norris et al., 1999), which can be observed in the admixture analysis where two ancestral populations were identified. The NOR population was established using fish from several rivers on the west coast in Norway, which probably favored greater genetic diversity than NAM and SCO, which showed just one ancestral population.

Recent events of admixture decrease the short-range LD present in original populations (Ødegård et al., 2014) and haplotypes with high LD levels are shorter in highly admixed populations (Toosi et al., 2010). Admixture can also generate long-range LD, which could be captured by lower density SNP panels (Ødegård et al., 2014; Vallejo et al., 2018). Conversely, the highest overall levels of LD present in the SCO and NAM populations may be reflecting the unique origin of these populations without the recent introgression of different genetic material and a small effective population size, as has been shown in Tasmanian Atlantic salmon (Kijas et al., 2017) and Pacific salmon (Barria et al., 2018).

Our results suggest that the effect of these demographic features is more extreme in the NAM population, which has the highest level of LD of the three populations analyzed. In general terms, LD varied moderately between chromosomes in the NAM population, suggesting a variation in autosomal recombination rate which could be associated to genetic drift or artificial selection (Arias et al., 2009). Using a MAF of 5% could results in biased estimates of population structure, as most structure-like clustering methods were not designed to account for the bias generated by MAF threshold (Guillot and Foll, 2009; Linck and Battey 2017). Therefore, our population structure analyses were performed without assessing for minor allele frequency quality control, allowing to include information from rare mutations along the different farmed populations.

The lower levels of SNP variability observed in the NAM population was expected, considering that North American salmon populations have lower genetic diversity than European populations (Bourret et al., 2013; Makinen et al., 2014). This also may be attributed to the ascertainment bias caused by prioritization of SNP markers segregating in NOR and SCO populations in the design of the SNP array used in the present study (Yáñez et al., 2016). A similar situation has been observed when evaluating performance SNP panels, which have been designed to account for the variability in European populations of Atlantic salmon and in Tasmanian farmed Atlantic salmon populations with a North American origin (Dominik et al., 2010; Kijas et al., 2017). To reduce this bias in the genetic differentiation analysis, we used a common subset of SNPs of approximately 33 K.

It has been suggested that the minimum number cattle needed for accurate LD estimations using r^2^ ranges between 55 and 75 individuals, increasing to more than 400 for |D’| (Bohmanova et al., 2010; Khatkar et al., 2008). However, an accurate estimation of LD measured as r^2^, has been obtained in Pacific salmon using 62 individuals (Barria et al., 2018). Because of the relatively small sample size of each population (71, 43 and 37 for the NOR, SCO and NAM populations, respectively), we measured LD decay as r^2^, instead of |D’|. Furthermore, estimates of r^2^ are less susceptible to overestimation and are more useful to predict the power of an association mapping (Ardlie et al., 2002; Bohmanova et al., 2010). Significant linear association has been assessed previously between chromosome length and LD (as r^2^) in Nellore cattle (Espigolan et al., 2013). We only found significance between these variables in the NOR population (p < 0.05). Like Bohmanova et al., 2010, we found no association in SCO and NAM populations, which could be due to lower marker density (data not shown).

The current results compared the LD decay between Chilean Atlantic salmon breeding populations originating from different geographic regions. The SNP panel used in the current study has one SNP every 14 Kb. Based on a r^2^ threshold value of 0.2 at a minimum marker distance of 42 kb, this panel can be used to detect associations between markers and traits of interest and also capture high-resolution information for genome predictions

### Conclusions

The current study reveals different LD decay between three Atlantic salmon farmed populations. The highest extent of LD was estimated for the NAM population, followed by the SCO and NOR populations. A lower level of LD in NOR was consistent with its higher level of admixture and consistent with population history. Specifically, this population comes from a farmed strain established with samples from several rivers in Norway. Therefore, subsequent genetic bottlenecks associated with strain formation have been less severe in comparison with the other two populations used in this study, that were established using fish from only one location. Also, the highest level of LD and lowest NE that was observed in NAM is consistent with the hypothesis that American salmon colonization from European fish favored only a few individuals. The high long range LD in NAM indicates the feasibility of achieving better prediction accuracies in this population with a smaller SNP data set than European populations.

### Ethics approval and consent to participate

The sampling protocol was previously approved by The Comité de Bioética Animal, Facultad de Ciencias Veterinarias y Pecuarias, Universidad de Chile (Certificate N° 29– 2014).

### Consent for publication

Not applicable

### Availability of data and material

The datasets used and/or analyzed during the current study are available from the corresponding author on reasonable request

## Author´s contributions

AB performed LD analysis and wrote the initial version of the manuscript. MEL performed population structure analysis, first quality control of genomic data and contributed with discussion and writing. GrY contributed with LD analysis and discussion. RC contributed with analysis and discussion. JMY conceived and designed the study, supervised work of AB and contributed to the analysis, discussion and writing. All authors have reviewed and approved the manuscript.

## Acknowledgements

AB acknowledge the National Commission of Scientific and Technologic Research (CONICYT) for the funding through the National PhD funding program.

## References

Ai, H., Huang, L., and Ren, J. (2013). Genetic Diversity, Linkage Disequilibrium and Selection Signatures in Chinese and Western Pigs Revealed by Genome-Wide SNP Markers. PLoS One 8.

Al-mamun, H. A., Clark, S. A., Kwan, P., and Gondro, C. (2015). Genome-wide linkage disequilibrium and genetic diversity in five populations of Australian domestic sheep. Genet. Sel. Evol. 47, 1–14.

Alexander, D., Novembre, J., and Lange, K. (2009). Fast Model-Based Estimation of Ancestry in Unrelated Individuals. Genome Res. 19, 1655–1664.

Ardlie, K. G., Kruglyak, L., and Seielstad, M. (2002). Patterns of Linkage Disequilibrium in the Human Genome. Nat. Rev. Genet. 3.

Arias, J. A., Keehan, M., Fisher, P., Coppieters, W., and Spelman, R. (2009). A high density linkage map of the bovine genome. BMC Genet. 10, 1–12.

Badke, Y. M., Bates, R. O., Ernst, C. W., Schwab, C., and Steibel, J. P. (2012). Estimation of linkage disequilibrium in four US pig breeds. BMC Genomics 13, 24.

Bangera, R., Correa, K., Lhorente, J. P., Figueroa, R., and Yáñez, J. M. (2017). Genomic predictions can accelerate selection for resistance against Piscirickettsia salmonis in Atlantic salmon (Salmo salar). BMC Genomics 18, 121.

Barbato, M., Orozco-terWengel, P., Tapio, M., and Bruford, M. W. (2015). SNeP: A tool to estimate trends in recent effective population size trajectories using genome-wide SNP data. Front. Genet. 6, 1–6.

Barria, A., Christensen, K. A., Yoshida, G., Jedlicki, A., Lhorente, J. P., Davidson, W. S., et al. (2018). Whole genome linkage disequilibrium and effective population size in a coho salmon (Oncorhynchus kisutch) breeding population. http://dx.doi.org/10.1101/335018.

Barría, A., Christensen, K. A., Yoshida, G. M., Correa, K., Jedlicki, A., Lhorente, J. P., et al. (2018). Genomic Predictions and Genome-Wide Association Study of Resistance Against Piscirickettsia salmonis in Coho Salmon (Oncorhynchus kisutch) Using ddRAD Sequencing. G3 Genes Genomes Genet. 8, 1183–1194.

Bohmanova, J., Sargolzaei, M., and Schenkel, F. S. (2010). Characteristics of linkage disequilibrium in North American Holsteins. BMC Genomics 11. doi:10.1186/1471-2164-11-421.

Bourret, V., Kent, M. P., Primmer, C. R., and Vasema, A. (2013). SNP-array reveals genome-wide patterns of geographical and potential adaptive divergence across the natural range of Atlantic salmon (Salmo salar). Mol. Ecol. 22, 532–551. doi:10.1111/mec.12003.

Corbin, L. J., Liu, A. Y. H., Bishop, S. C., and Woolliams, J. A. (2012). Estimation of historical effective population size using linkage disequilibria with marker data. J. Anim. Breed. Genet. 129, 257–270.

Correa, K., Bangera, R., Figueroa, R., Lhorente, J. P., and Yáñez, J. M. (2017). The use of genomic information increases the accuracy of breeding value predictions for sea louse (Caligus rogercresseyi) resistance in Atlantic salmon (Salmo salar). Genet. Sel. Evol. 49, 15.

Correa, K., Lhorente, J. P., López, M. E., Bassini, L., Naswa, S., Deeb, N., et al. (2015). Genome-wide association analysis reveals loci associated with resistance against Piscirickettsia salmonis in two Atlantic salmon (Salmo salar L.) chromosomes. BMC Genomics 16, 854.

Dominik, S., Henshall, J. M., Kube, P. D., King, H., Lien, S., Kent, M. P., et al. (2010). Evaluation of an Atlantic salmon SNP chip as a genomic tool for the application in a Tasmanian Atlantic salmon (Salmo salar) breeding population. Aquaculture 308, S56–S61. doi:10.1016/j.aquaculture.2010.05.038.

Espigolan, R., Baldi, F., Boligon, A. A., Souza, F. R. P., Gordo, D. G. M., Tonussi, R. L., et al. (2013). Study of whole genome linkage disequilibrium in Nellore cattle. BMC Genomics 14.

FAO (2016a). Fisheries and Aquaculture Information and Statistical Branch.

FAO (2016b). The state of world fisheries and aquaculture.

Flint-Garcia, S. A., Thornsberry, J. M., and Buckler, E. S. (2003). Structure of Linkage Disequilibrium in Plants. Annu. Rev. Plant Biol. 54, 357–374.

Gaut, B. S., and Long, A. D. (2003). The Lowdown on Linkage Disequilibrium. Plant Cell 15, 1502–1507.

Gjedrem, T., Robinson, N., and Rye, M. (2012). The importance of selective breeding in aquaculture to meet future demands for animal protein: A review. Aquaculture 350–353, 117–129.

Goddard, M., and Hayes, B. (2009). Mapping genes for complex traits in domestic animals and their use in breeding programmes. Nat. Rev. Genet. 10, 381–391.

Guillot, G., and Foll, M. (2009). Correcting for ascertainment bias in the inference of population structure. Bioinformatics 25, 552–554.

Gutierrez, A. P., Yáñez, J. M., Fukui, S., Swift, B., and Davidson, W. S. (2015). Genome-Wide Association Study (GWAS) for Growth Rate and Age at Sexual Maturation in Atlantic Salmon (Salmo salar). PLoS One 10, e0119730.

Hill, W., and Robertson, A. (1968). Linkage disequilibrium in finite populations. Theor. Appl. Genet. 38, 226–231.

Houston, R. D., Taggart, J. B., Cézard, T., Bekaert, M., Lowe, N. R., Downing, A., et al. (2014). Development and validation of a high density SNP genotyping array for Atlantic salmon (Salmo salar). BMC Genomics 15, 90.

Johnston, I. a, McLay, H. a, Abercromby, M., and Robins, D. (2000). Phenotypic plasticity of early myogenesis and satellite cell numbers in atlantic salmon spawning in upland and lowland tributaries of a river system. J. Exp. Biol. 203, 2539–2552.

Khatkar, M. S., Nicholas, F. W., Collins, A. R., Zenger, K. R., Cavanagh, J. A. L., Barris, W., et al. (2008). Extent of genome-wide linkage disequilibrium in Australian Holstein-Friesian cattle based on a high-density SNP panel. BMC Genomics 9.

Kijas, J., Elliot, N., Kube, P., Evans, B., Botwright, N., King, H., et al. (2017). Diversity and linkage disequilibrium in farmed Tasmanian Atlantic salmon. Anim. Genet. 48, 237–241.

Linck, E., and Battey, C. (2017). Minor allele frequency thresholds strongly affect population structure inference with genomic datasets. bioRxiv, https://doi.org/10.1101/188623.

López, M. E., Benestan, L., Moore, J.-S., Perrier, C., Gilbey, J., Di Genova, A., et al. (2018). Comparing genomic signatures of domestication in two Atlantic salmon (Salmo salar L) populations with different geographical origins. Evol. Appl., 0–2.

Lu, D., Sargolzaei, M., Kelly, M., Li, C., Voort, G. Vander, Wang, Z., et al. (2012). Linkage disequilibrium in Angus, Charolais, and Crossbred beef cattle. Front. Genet. 3, 1–10.

Makina, S. O., Taylor, J. F., Van Marle-Köster, E., Muchadeyi, F. C., Makgahlela, M. L., MacNeil, M. D., et al. (2015). Extent of linkage disequilibrium and effective population size in four South African sanga cattle breeds. Front. Genet. 6, 1–12.

Makinen, H., Vasemagi, A., Mcginnity, P., Cross, T. F., and Primmer, C. R. (2014). Population genomic analyses of early-phase Atlantic Salmon (Salmo salar) domestication/captive breeding. Evol. Appl. 8, 93–107. doi:10.1111/eva.12230.

McKay, S. D., Schnabel, R. D., Murdoch, B. M., Matukumalli, L. K., Aerts, J., Coppieters, W., et al. (2007). Whole genome linkage disequilibrium maps in cattle. BMC Genet. 8, 74.

Mdladla, K., Dzomba, E. F., Huson, H. J., and Muchadeyi, F. C. (2016). Population genomic structure and linkage disequilibrium analysis of South African goat breeds using genome-wide SNP data. Anim. Genet. 47, 471–482.

Meuwissen, T. H., Hayes, B. J., and Goddard, M. E. (2001). Prediction of Total Genetic Value Using Genome-Wide Dense Marker Maps. Genetics 157, 1819–1829.

Norris, A. T., Bradley, D. G., and Cunningham, E. P. (1999). Microsatellite genetic variation between and within farmed and wild Atlantic salmon (Salmo salar) populations. Aquaculture 180, 247–264.

Ødegård, J., Moen, T., Santi, N., Korsvoll, S. A., Kjøglum, S., and Meuwisse, T. H. E. (2014). Genomic prediction in an admixed population of Atlantic salmon (Salmo salar). Front. Genet. 5, 1–8.

Porto-Neto, L. R., Kijas, J. W., and Reverter, A. (2014). The extent of linkage disequilibrium in beef cattle breeds using high-density SNP genotypes. Genet. Sel. Evol. 46, 1–5.

Prasad, A., Schnabel, R. D., McKay, S. D., Murdoch, B., Stothard, P., Kolbehdari, D., et al. (2008). Linkage disequilibrium and signatures of selection on chromosomes 19 and 29 in beef and dairy cattle. Anim. Genet. 39, 597–605.

Prieur, V., Clarke, S. M., Brito, L. F., McEwan, J. C., Lee, M. A., Brauning, R., et al. (2017). Estimation of linkage disequilibrium and effective population size in New Zealand sheep using three different methods to create genetic maps. BMC Genet. 18, 1–19.

Pritchard, J. K., and Przeworski, M. (2001). Linkage Disequilibrium in Humans: Models and Data. Am. J. Hum. Genet. 69, 1–14. doi:10.1086/321275.

Purcell, S., Neale, B., Todd-Brown, K., Thomas, L., Ferreira, M. A. R., Bender, D., et al. (2007). PLINK: A tool set for whole-genome association and population-based linkage analyses. Am. J. Hum. Genet. 81, 559–575.

R Core Team (2016). R: A Language and Environment for Statistical Computing; R Foundation for Statistical Computing: Vienna, Austria, 2015.

Rexroad, C. E., and Vallejo, R. L. (2009). Estimates of linkage disequilibrium and effective population size in rainbow trout. BMC Genet. 10, 83.

Rougemont, Q., and Bernatchez, L. (2018). The demographic history of Atlantic Salmon (Salmo salar) across its distribution range reconstructed from Approximate Bayesian Computations. Evolution, https://doi.org/10.1111/evo.13486.

Rye, M., Gjerde, B., and Gjedrem, T. (2010). Genetic Improvement Programs For Aquaculture Species In Developed Countries. in World Congress on Genetics Applied to Livestock Production.

Solar, I. I. (2009). Use and exchange of salmonid genetic resources relevant for food and aquaculture. Rev. Aquac. 1, 174–196.

Sonesson, A. K., and Meuwissen, T. H. E. (2009). Testing strategies for genomic selection in aquaculture breeding programs. Genet. Sel. Evol. 41, 37.

Sved, J. A. (1971). Linkage disequilibrium and homozygosity of chromosome segments in finite populations. Theor. Popul. Biol. 2, 125–141.

Toosi, A., Fernando, R. L., and Dekkers, J. C. M. (2010). Genomic selection in admixed and crossbred populations. J. Anim. Sci. 88, 32–46.

Tsai, H. Y., Hamilton, A., Tinch, A. E., Guy, D. R., Bron, J. E., Taggart, J. B., et al. (2016). Genomic prediction of host resistance to sea lice in farmed Atlantic salmon populations. Genet. Sel. Evol. 48, 1–11.

Vallejo, R. L., Leeds, T. D., Gao, G., Parsons, J. E., Martin, K. E., Evenhuis, J. P., et al. (2017). Genomic selection models double the accuracy of predicted breeding values for bacterial cold water disease resistance compared to a traditional pedigree-based model in rainbow trout aquaculture. Genet. Sel. Evol. 49, 17.

Vallejo, R. L., Silva, R. M. O., Evenhuis, J. P., Gao, G., Sixin, L., Parsons, J. E., et al. (2018). Accurate genomic predictions for BCWD resistance in rainbow trout are achieved using low-density SNP panels?: Evidence that long-range LD is a major contributing factor. J. Anim. Breed. Genet. doi:10.1111/jbg.12335.

Verspoor, E., Stradmeyer, L., and Nielsen, J. L. (2007). The Atlantic Salmon: Genetics, Conservation and Management.

Visser, C., Lashmar, S. F., Marle-köster, E. Van, Poli, M. A., and Allain, D. (2016). Genetic Diversity and Population Structure in South African, French and Argentinian Angora Goats from Genome-Wide SNP Data. PLoS One 11, e0154353.

Yang, J., Shikano, T., Li, M., and Merilä, J. (2014). Genome-Wide Linkage Disequilibrium in Nine-Spined Stickleback Populations. G3 4, 1919–1929.

Yáñez, J. M., Bangera, R., Lhorente, J. P., Oyarzún, M., and Neira, R. (2013). Quantitative genetic variation of resistance against Piscirickettsia salmonis in Atlantic salmon (Salmo salar). Aquaculture 414–415, 155–159.

Yáñez, J. M., Lhorente, J. P., Bassini, L. N., Oyarzún, M., Neira, R., and Newman, S. (2014). Genetic co-variation between resistance against both Caligus rogercresseyi and Piscirickettsia salmonis, and body weight in Atlantic salmon (Salmo salar). Aquaculture 433, 295–298. doi:10.1016/j.aquaculture.2014.06.026.

Yáñez, J. M., Naswa, S., Lopez, M. E., Bassini, L., Correa, K., Gilbey, J., et al. (2016). Genomewide single nucleotide polymorphism discovery in Atlantic salmon (Salmo salar): validation in wild and farmed American and European populations. Mol. Ecol. Resour. 16, 1002–1011.

Yáñez, J. M., Scott, N., and Houston, R. D. (2015). Genomics in aquaculture to better understand species biology and accelerate genetic progress. Front. Genet. 6, 1–3.

Yoshida, G. M., Bangera, R., Carvalheiro, R., Correa, K., Figueroa, R., Lhorente, J. P., et al. (2018). Genomic Prediction Accuracy for Resistance Against Piscirickettsia salmonis in Farmed Rainbow Trout. G3 Genes Genomes Genet. 8, 719–726. doi:10.1534/g3.117.300499.

Zhao, H., Nettleton, D., Zoller, M., and Dekkers, J. C. M. (2005). Evaluation of linkage disequilibrium measures between multi-allelic markers as predictors of linkage disequilibrium between markers and QTL. Genet. Res. 86, 77–87.

